# A comprehensive analysis of Usutu virus (USUV) genomes revealed lineage-specific codon usage patterns and host adaptation

**DOI:** 10.1101/2022.05.04.490698

**Authors:** Jianglin Zhou, Yaling Xing, Zhe Zhou, Shengqi Wang

**Affiliations:** Beijing institute of microbiology and epidemiology, Beijing 100850, China

**Keywords:** codon usage, natural selection, mutation pressure, evolution, Usutu virus

## Abstract

The Usutu virus (USUV) is an emerging arbovirus virus maintained in the environment of Afro-Eurasia via a bird-mosquito-bird enzootic cycle and sporadically infected other vertebrates. Despite primarily asymptomatic or mild symptoms, humans infected by USUV can develop severe neurological diseases such as meningoencephalitis. However, no detailed study has yet been conducted to investigate its evolution from the perspective of codon usage patterns. Codon usage choice of viruses reflects the genetic variations that enable them to reconcile their viability and fitness towards the external environment and new hosts. This study performed a comprehensive evolution and codon usage analysis in USUVs. Our reconstructed phylogenetic tree confirmed the circulation viruses belonging to eight distinct lineages, reaffirmed by principal component analysis based on codon usage patterns. We also found a relatively small codon usage bias and that natural selection, mutation pressure, and evolutionary processes collectively shaped the codon usage of the USUV, with natural selection predominating over the others. Additionally, a complex interaction of codon usage between the USUV and its host was observed. This process could have enabled USUVs to adapt to various hosts and vectors, including humans. Therefore, the USUV may possess a potential risk of cross-species transmission and subsequent outbreaks. In this respect, further epidemiologic surveys, diversity monitoring, and pathogenetic research are warranted.

## Introduction

Usutu virus (USUV) is an emerging arbovirus belonging to the genus *Flavivirus* in the family *Flaviviridae*. USUV is a member of the Japanese encephalitis virus (JEV) serocomplex, genetically close to human pathogens JEV, West Nile virus (WNV), and Murray Valley encephalitis virus (MVEV) [1]. Like other flaviviruses, USUV has a38 +ss RNA genome comprised of 11064 nucleotides that encodes one open reading frame (ORF) and two flanking untranslated regions[2,3]. The ORF that encodes a polyprotein of 3434 amino acids will be enzymatically cleaved into three structural proteins (C, prM, E) and seven non-structural proteins (NS1, NS2A, NS2B, NS3,NS4A, NS4B, and NS5). Since first isolated in 1959 from a *Culex neavei* mosquito in Swaziland, USUV has continuously circulated within Africa and later spread across Europe [4]. USUV is sustained in an enzootic cycle among wild birds as amplifying hosts (primarily in *Turdus merula*) and mosquitoes as vectors (mainly in *Cx. pipiens*).Humans and other mammals, including rodents, horses, bats, and deer, are sporadically infected and considered to be dead-end hosts[1,4]. To date, at least 100cases of acute infection have been described in humans, with symptoms ranging from mild or asymptomatic to severe neurological disease [5]. Besides epidemic potential, USUV may also represent a risk for blood safety, especially in the context of co-circulation with WNV and probably underestimated circulation of USUV [6,7].Therefore, an exhaustive study of the replication and evolution of USUVs is warranted.

The redundancy of genetic code allows organisms to regulate their efficiency and accuracy of protein production while preserving the same amino-acid sequences [8,9]. During protein translation in a certain specie or cell, some codons are used more frequently than others, a phenomenon is known as codon usage bias (CUB) [10,11]. Previous studies indicated that CUB is common in three domains and viruses and is influenced by many factors, such as mutation pressure, natural or translation selection, dinucleotide abundance, and external environment [8,12-14]. Considering the entire parasitism of viruses, the interactions of the virus and its host are expected to influence viral viability, fitness, evolution, and evasion of the host immune responses [8,13]. Studying CUB thus supplies a novel perspective on virus evolution and can deepen our understanding of the biological properties of USUVs and aid in potential vaccine design. However, to our knowledge, there is only one report on the codon adaptation index for just four hosts within a fraction of USUVs [15]; no detailed analysis of codon usage of USUVs has been published.

In this study, we comprehensively analyzed the phylogenetic relationships and codon usage patterns of USUVs. We also explored the possible key factors responsible for the CUB of USUV as well as its adaptation to various hosts. Our results show a novel perspective regarding the molecular evolution in USUV.

## Materials and methods

### Dataset retrieval and annotation

All whole genomes of USUV were collected from the GenBank database on March 10, 2022. One sequence was kept for identical sequences. The USUV genomes were annotated by VADR [16]. Genomes whose ORF has fuzzy coordinate or non-(A, C, G, U) nucleotides were removed. Finally, a total of 368 genomes were analyzed in this study. Detailed genomes are listed in Table S1.

### Recombination and phylogenetic analysis

Potential recombination events in USUV coding sequences were detected by the Genetic Algorithm for Recombination Detection (GARD) using Datamonkey web service [17,18]. All genome sequences were aligned by MAFFT [19]. The maximum likelihood (ML) phylogenetic tree was constructed by IQ-TREE with 1000 replications of ultrafast bootstrap resampling [20] and SH-aLRT test [21]. The model GTR+F+I+G4 was selected using the built-in ModelFinder [22]. The tree was visualized using the ggtree package [23].

### Nucleotide and codon composition analysis

The frequencies of A, U, G, and C, overall GC content, GC percentage at the first (GC1s), second (GC2s), third (GC3s) codon position and the average of GC1s and GC2s (GC12s) were calculated by seqinr package [24]. The frequencies of A, U, G, and C at the third positions in the synonymous codons (A3s, U3s, G3s, C3s) were calculated by CodonW (http://codonw.sourceforge.net/culong.html#CodonW). Five codons without synonymous codons, including AUG, UGG, UAG, UAA, and UAG, were excluded from this analysis.

### Relative synonymous codon usage (RSCU) analysis

The RSCU values represent the usage frequencies of synonymous codons in protein excluding the effect of the sequence length and amino acid compositions[25]. The RSCU value was estimated using the seqinr package as follows:

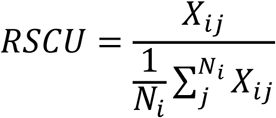

Where X*ij* is the observed number of *j*th codon for the *i*th amino acid, which has *N*_*i*_ kinds of alternative synonymous codons. Codons with RSCU values > 1.6 are considered as over-represented, whereas < 0.6 reflected under-represented ones.

### Principal component analysis (PCA)

PCA is widely used to resolve the relationship between the multivariate and samples. Here, each ORF represented by a 59-dimensional vector was transformed into several principal components (PCs). The PCA analysis was performed using the factoextra package [26].

### The effective number of codons (ENC) estimation

The ENC value indicates the extent of CUB, ranging from 20 to 61 [27]. The smaller ENC value represents a stronger CUB. The ENC values were estimated by the seqinr package。

### ENC-plot analysis

To identify factors influencing CUB, ENC-plot analysis was performed by plotting the ENC values against the GC3s. Genes whose codon usage is only constrained by mutation pressure will locate on or around the expected curve. Otherwise, natural selection exerts a more powerful influence. The expected ENC value was inferred using the below formula:

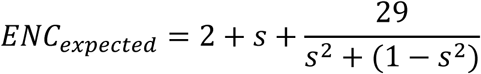

Where the *s* represents values of GC3s [28].

### Neutrality plot analysis

The neutrality plot was used to determine the dominant factors (mutation pressure or natural selection) influencing CUB [29]. The GC12s values (y-axis) were plotted against the GC3s values (x-axis). Mutation pressure is considered the dominant force shaping codon usage if the coefficient of GC3s is statistically significant and close to 1. The slope value is closer to 0 means a higher influence from natural selection.

### Codon adaption index (CAI) calculation

The CAI analysis is a quantitative method that is applied to evaluate the adaptiveness of a gene toward the codons of highly expressed genes [30]. CAI values of USUV coding sequences were calculated using the local version of CAIcal [31], to the codon usage patterns of its hosts and vectors. A total of 14 species representing four categories of hosts and vectors, including birds (*T. merula, Sturnus vulgaris, Passer domesticus, Alauda arvensis*, and *Bubo scandiacus*), mosquitoes (*Cx. pipiens* pallens, *Cx. quinquefasciatus, Aedes albopictus*, and *Ae. aegypti*), human (*Homo sapiens*) and non-human mammals (NHM, *Pipistrellus pipistrellus, Rattus rattus, Capreolus capreolus*, and *Equus caballus*), were obtained from the Codon and Codon-Pair Usage Tables (CoCoPUTs) database on March 21, 2022 [32]. The CAI value by reference codon usage pattern ranges from 0 to 1 with higher CAI values signifying better viral adaptation to the corresponding host.

### Relative codon deoptimization index (RCDI) calculation

The RCDI value measures the deoptimization of the USUV towards that of its hosts. An RCDI value of 1 indicates that the virus pursues the codon usage pattern of the host and exhibits a host-adapted codon usage preference. Contrarily, an RCDI value >1 indicates the codon usage pattern of the virus deviates from its host. The RCDI values were calculated for the 14 species using CAIcal [31].

### Similarity index analysis

The similarity index [SiD or D(A, B)] is an indicator to estimate the overall effect of host codon usage on viral codon usage. The SiD values of USUV for fourteen hosts were calculated using the following formula [33]:

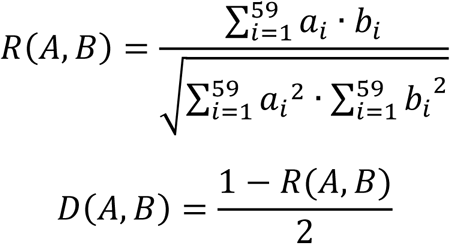

Where *a*_*i*_ and *b*_*i*_ represent the RSCU value of 59 synonymous codons for the USUV and the host, respectively. *D(A, B)* indicates the potential effect of the overall codon usage of the host on that of USUV, ranging from 0 to 1. A Higher SiD value means a greater impact from the host on USUV codon usage.

### Correlation and statistics analysis

Spearman’s rank correlation analysis was performed to determine the relationships among the genomic composition, ENC, Aromo, Gravy, and the first two axes of PCA. A two-sided Dunn’s test was used to the statistical significance between groups. *P* values were corrected using Benjamini-Hochberg (BH) procedure and 0.05 was used as the significance threshold.

## Results

### Phylogenetic analysis of USUV

GARD analysis found no evidence of recombination event among the 368 USUV strains, hence all of them were included for subsequent phylogenetic and codon usage analyses. The obtained ML phylogeny shows that these USUV strains fell into eight distinct African (AF) and European (EU) lineages, namely AF1-3, and EU1-5 (Figure 1). The AF1, which contains only one strain from an African mosquito, is distantly far away from the others. Except for the EU4, which only includes viruses from Italy, AF2, AF3, EU1, EU2, EU3 and EU5 are widespread in varied hosts of many countries and continents.

**Figure 1.**
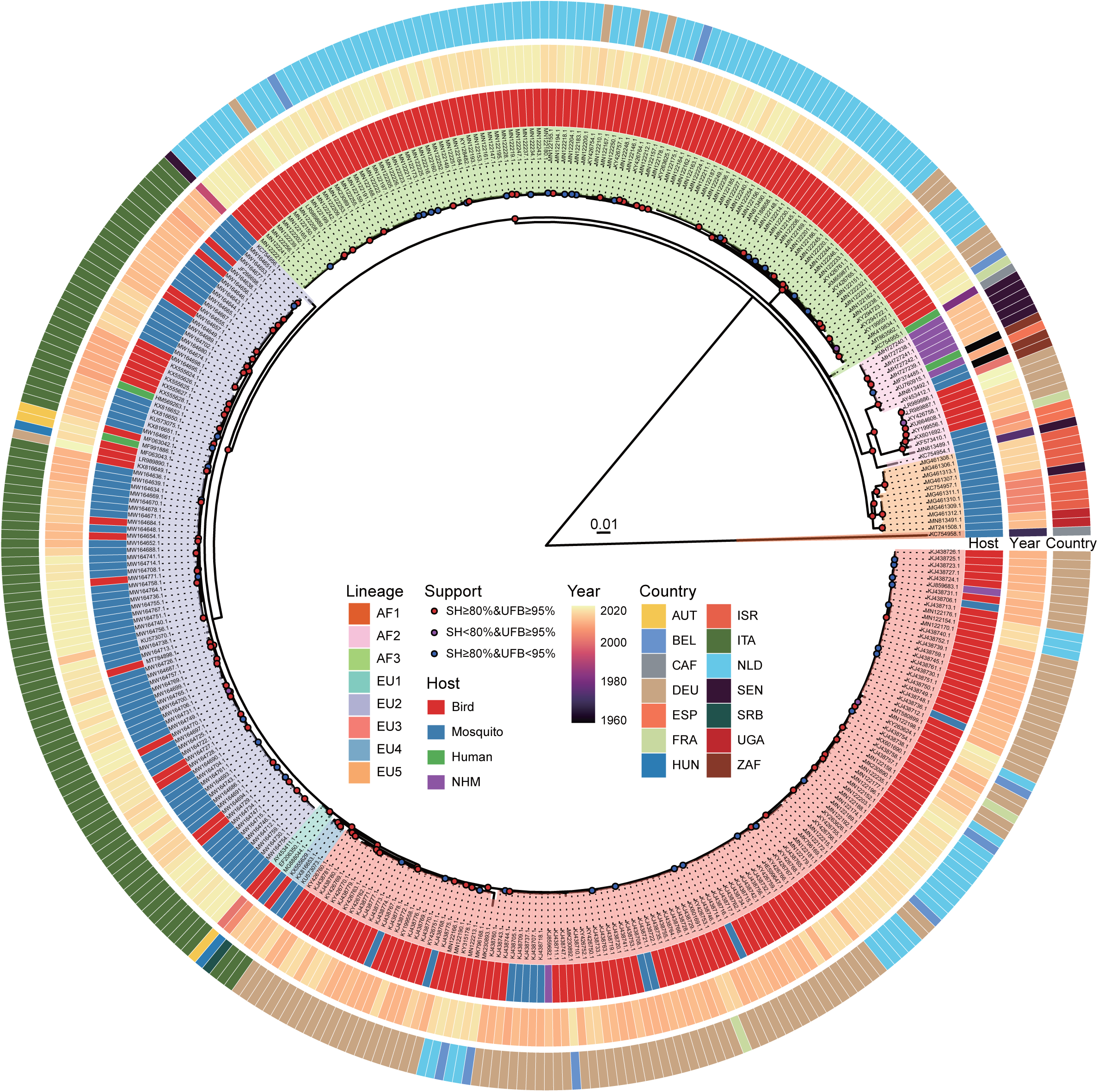
Phylogenetic tree of 368 whole genomes of USUV based on IQ-TREE. The background of the USUV strains labels was filled based on lineage classification.

### G and A nucleotides are more abundant in the USUV coding sequences

The nucleotide composition was analyzed to evaluate its potential impact on codon usage of USUV. Here we found that the most frequent mononucleotide was G, with a mean ± standard deviation (SD) value of 28.34 ± 0.08%, followed by A (27.03 ±176 0.08%), C (22.87 ± 0.07%), and U (21.76 ± 0.08%). The C3s, A3s, G3s, and U3s was 177 34.07 ± 0.22%, 30.86 ± 0.28%, 30.71 ± 0.26%, and 26.48 ± 0.20%, respectively. The178 overall GC content (51.21 ± 0.06%) was slightly higher than that of AU. The GC1s 179 (56.91 ± 0.09%) and GC3s (52.34 ± 0.17%) values were higher than GC2s (44.91 ±0.05%) and GC12s (50.91 ± 0.06%). The detailed nucleotide compositions of strains are listed in Table S2. Therefore, although the USUV coding sequences were GC-rich, mononucleotides G and A were more abundant. Significant differences (adjusted *P* <0.05) were also noticed in the average GC, GC1s, GC2s, and GC3s values of USUV strain in various lineages and hosts (FigS1). These results confirmed that nucleotide compositions of the USUV viruses are complicated and imbalanced, implying a biased codon usage.

### CUB among the USUV

The ENC values were calculated to estimate the degree of USUV CUB. The ENC values of whole coding sequences ranged from 54.95 to 56.05 (mean 55.30 ± 0.19),irrespective of lineage (Table S2). Concerning the lineage classification of complete coding sequences, a significantly highest ENC value of 55.60 ± 0.33 was observed in the AF2 lineage while the lowest ENC value of 55.08 ± 0.05 was observed in the EU3 lineage (*P* <0.0001, Figure 2A). Analyzing individual genes showed the ENC values of individual genes of different lineages exhibited distinguishing characteristics, especially the AF1 lineage (Figure 2B). Significant disparities (adjusted *P* < 0.05)were discovered in the average ENC values of the ten genes and different lineages of each gene (FigS2). These results suggested a low and lineage-specific CUB among the USUV coding sequences.

**Figure 2.**
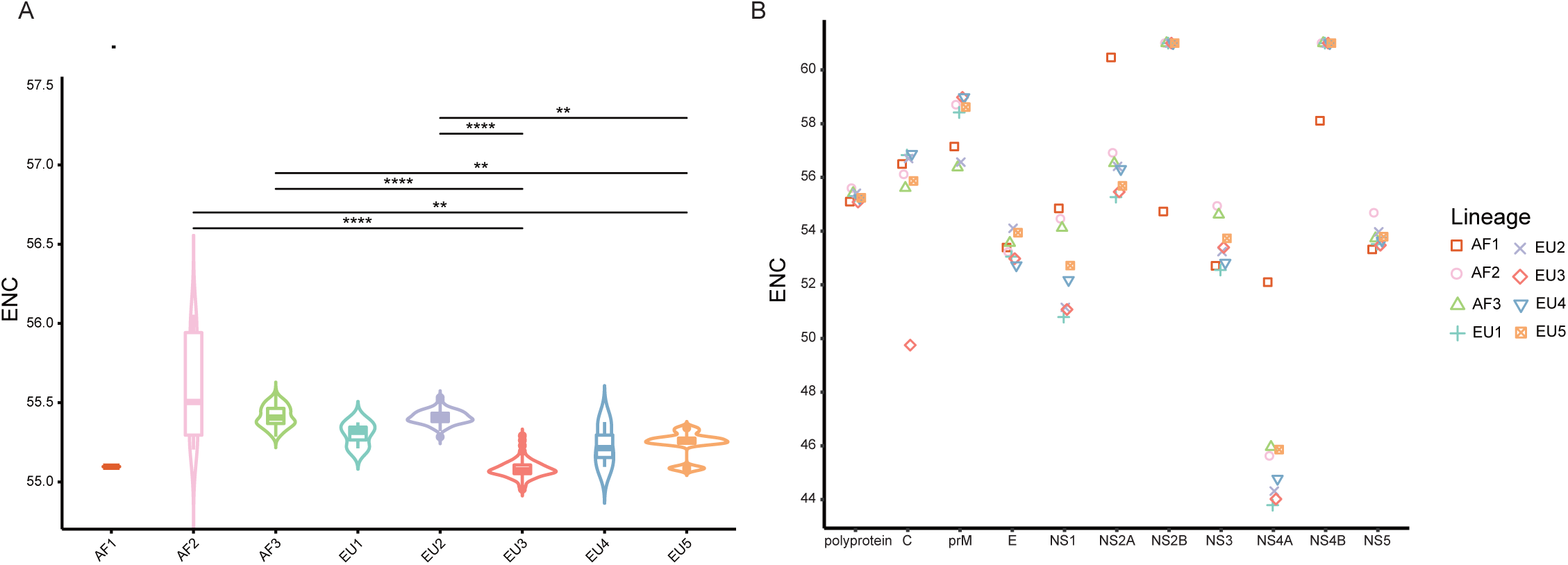
The ENC values distribution. (A) The violin plots with inner boxplots showed the ENC values of polyproteins of USUV in different lineages. All differences with *P* < 0.05 are indicated. \*\**P* < 0.01; \*\*\**P* < 0.001; \*\*\**P* < 0.0001. (B) The scatter plot of ENC values of the various gene from different lineages.

### USUVs have evolved into lineage-specific RSCU patterns

RSCU analysis is used to explore the patterns of and preferences for codon usage among genes. Here we found that except for Phe without CUB, all the remaining 17 amino acids had preferred codons (RSCU > 1.0) (TableS3). Specifically, 29 of 59 synonymous codons were classified as preferred codons, eighteen of them are G/C-ended (12 C-ended; 6 G-ended) and eleven were A/U-ended (7 A-ended; 4 U-ended).This means C- and A-ended codons are preferred in the USUV. Among the preferred codons, three codons (AGA, CUG, GGA) were over-represented (RSCU > 1.6).Similarly, nine codons (UUA, GUA, UCG, CCG, ACG, GCG, CGA, CGC, CGU)were under-represented (RSCU < 0.6).

Taking lineage information into consideration, we found that preferred codons varied. A total of 35 codons were preferred by at least one lineage, while only 21 of them were preferred by all eight lineages. The preferred codons of some amino acids were different among the lineages (TableS3). Moreover, unlike the other lineages, the AF1 lineage had five over-represented codons and three of them are unique (GUG,CCA, and AGG). The underrepresentation analysis result was more complex. A total of 11 codons were under-represented in at least one lineage, and 6 of them were under-represented in all eight lineages. The heatmap also indicated distinctively lineage-specific RSCU patterns (Figure 3). The lineage-specific codon usage patterns underscore the independent evolutionary history of USUV strains. Additionally, we found that the common preferred codons (RSCU > 1.0) and unpreferred codons (RSCU < 1.0) were neither completely harmonious nor opposite in USUV compared to any of the hosts (TableS3).

**Figure 3.**
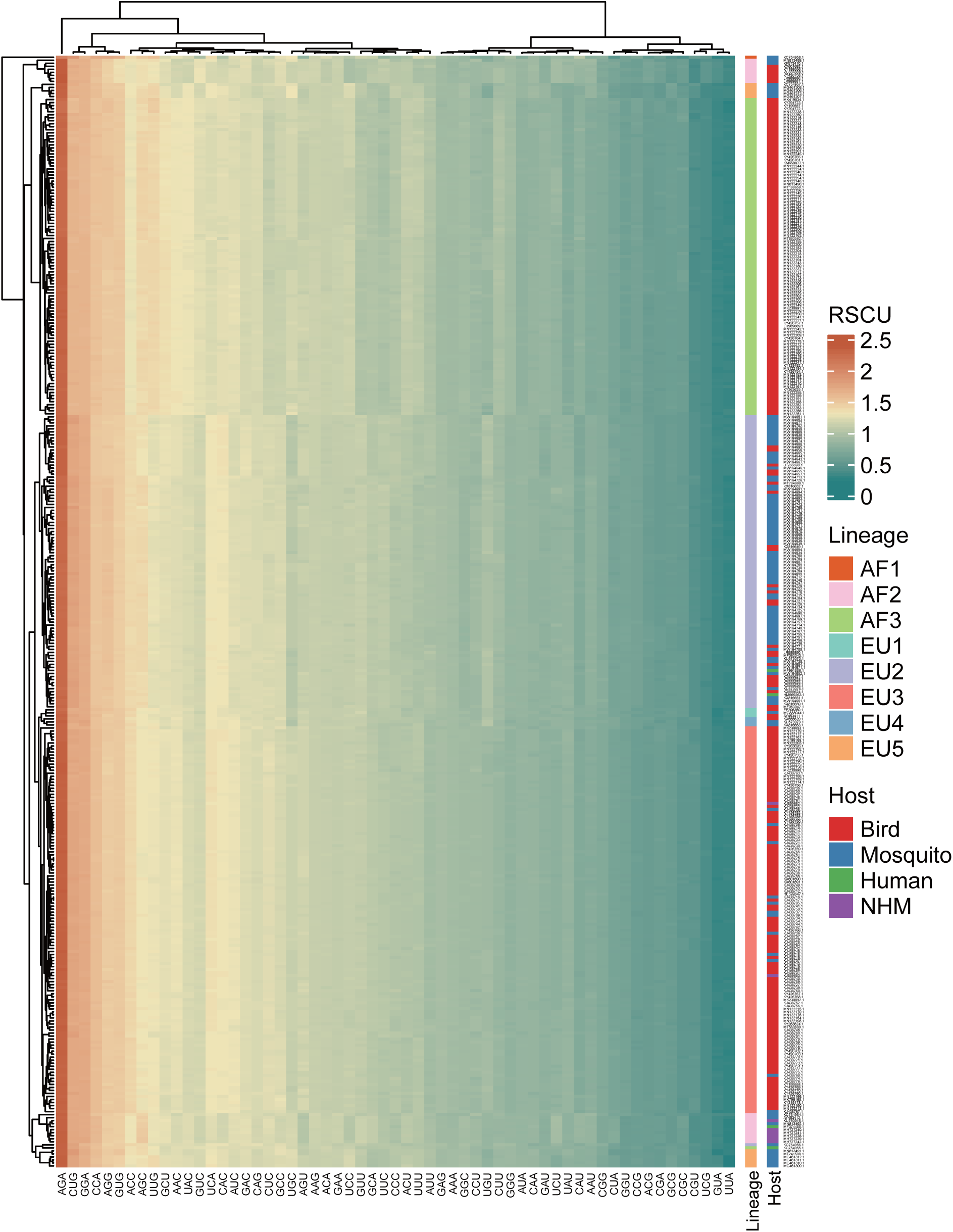
Heatmap of mean RSCU values among the 368 complete coding sequences of USUV. Each row represents a USUV isolate and each column represents a codon.

### Trends of codon usage variations in USUV

PCA analysis was performed to explore the synonymous codon usage variations among the USUV isolates. The first and second axes accounted for 41.08% and 14.12% of the total codon usage variation (Figure 4A). The strains were mainly grouped into five well-defined clusters, corresponding to 5 of 8 lineages (AF2, AF3,EU2, EU3, and EU5). The remaining three lineages were scattered probably due to their small population size. Specifically, the AF2, AF3, EU2, and EU3 lineages were grouped into distinctly separate clusters. However, the 95% confidence ellipse of the EU5 lineage had a few overlapping with that of the AF2 and AF3 lineages. The AF1 didn’t closely group with any clusters/lineages. In addition, we also performed PCA of individual genes based on lineages and host (Figure 4B). The unique codon usage of the AF1 lineage is retained in all genes. Instead, the distinctly separated clusters of the five lineages were kept in some genes such as *E* and *NS5*, whereas much more overlapping tendencies were found in the other genes such as *C* and *NS2B*. All above, these results reconfirm a lineage-specific codon usage of USUV and suggest a common ancestry, but the independent and varied divergence history at the levels of individual genes.

**Figure 4.**
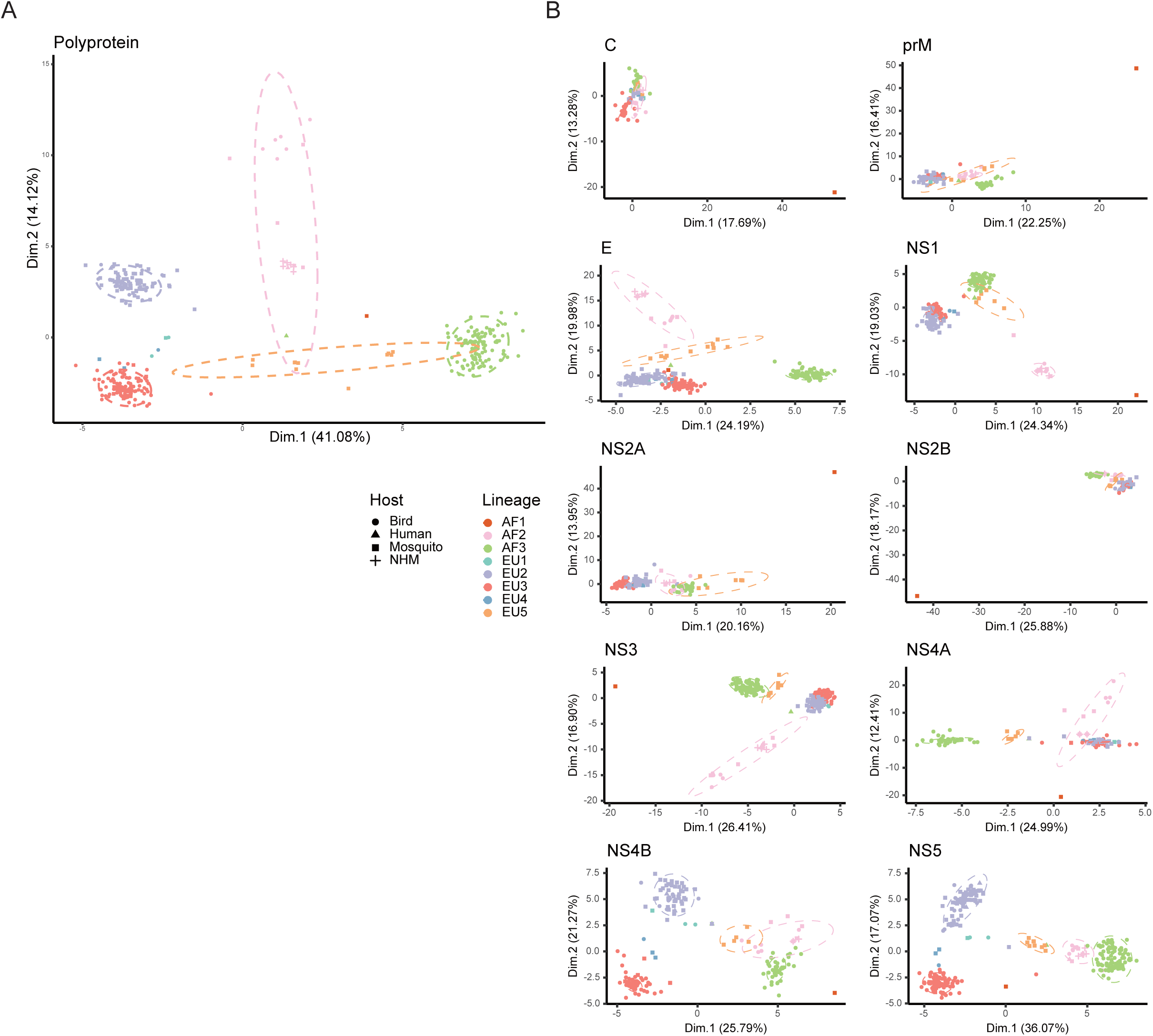
PCA based on the RSCU values of 59 synonymous codons. PCA biplots were performed on whole coding sequences (A) and every gene (B). The ellipses are drawn in a 95% confidence interval.

### Both natural selection and mutation pressure shape the codon usage pattern of USUV

To determine the factors that influence the codon usage pattern, ENC plots and correlation analyses were performed. In the ENC-GC3s plot of complete coding sequences, all USUV isolates were lying below the expected ENC curve (Figure 5).The strains from AF2, EU3, AF3, EU2, and EU5 formed five distinguishing clusters, albeit clusters of the later three lineages had a few overlaps. This indicated natural selection dominated the codon usage of all USUV strains. However, ENC plots of individual genes showed that the effects of natural selection and mutation pressure on codon usage varied. For example, all *NS2B* and *NS4B* coding sequences sat above the expected ENC curve, except for the AF1, showing the dominant role of mutation pressure in these genes (Figure S3). These results suggested that both mutation and natural selection shape the codon usage patterns of USUVs. Furthermore, correlation analysis revealed a mixture of significant (*P* < 0.05)and non-significant correlations between nucleotide compositions and codon compositions (Figure S4). Especially, the first two axes of PCA had significant correlations with almost all the indices, including mononucleotides, A3s, C3s, G3s,U3s, GC1s, Aromo, and ENC. A remarkable relationship between mononucleotides, A3s, C3s, G3s, U3s, and ENC was observed as well (all |*r*| ≥ 0.63). These results reconfirmed the combined role of mutation pressure and natural selection in the codon usage propensities of USUV.

**Figure 5.**
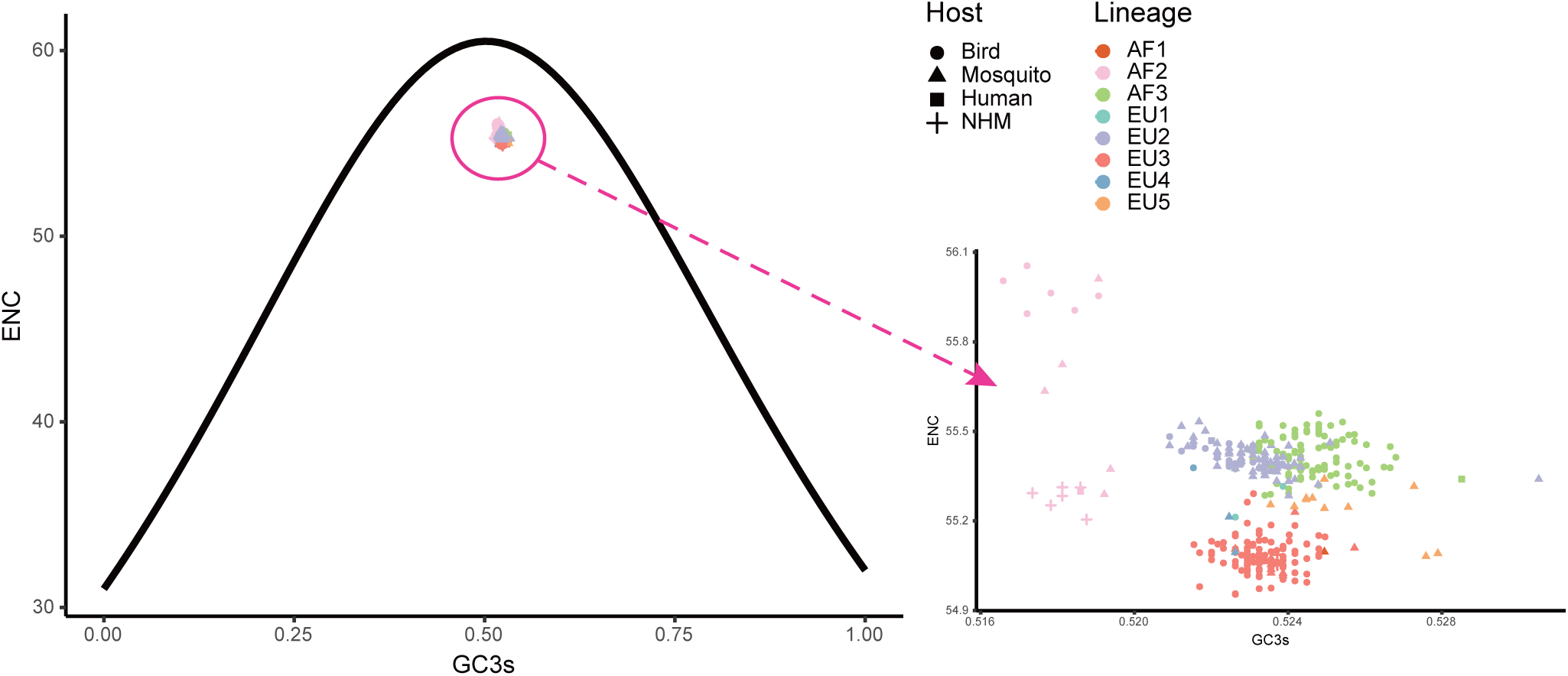
The ENC plot of whole coding sequences of USUVs. The solid curve represents the expected ENC values when the codon usage was only influenced by the GC3s composition.

### Natural selection is the major driver of USUV codon usage

Once we recognized that both natural selection and mutation pressure contributed to the CUB of the USUV, a neutrality analysis was conducted to determine the magnitude of the two forces. Regarding complete coding sequences, neutrality analysis showed a low but significant correlation between GC12s and GC3s values among all the strains (R^2^_adj_ = 0.069, *P* < 0.0001). The slope of the regression line was inferred to be -0.09, according to which mutation pressure (relative neutrality) was 9% and natural selection (the relative constraint on GC3s) was 91% (Figure 6A), indicating the principal effect of natural selection on the codon usage of USUV. Based on individual lineages analyses, the slopes of linear regression were 0.36, 0.00092, -0.5, -0.023, 0.015, 0.51, and -0.12 for the AF2-3 and EU1-5 lineages, respectively (Figure 6B). Therefore, the mutation pressure accounted for 36%, 0.092%, 50%, 2.3%, 1.5%, 51% and 12%, whereas natural selection accounted for 64%, 99.908%, 50%, 97.7%, 98.5%, 49% and 88% in the corresponding lineages, respectively. The AF1 lineage had no linear regression result due to its single population size. Although mutation pressure explained 50% and 51% in the EU1 and EU4 lineage, respectively, all the correlations were not statistically significant in the seven lineages (*P* > 0.2392). These results reaffirmed the dominant influence of natural selection.

**Figure 6.**
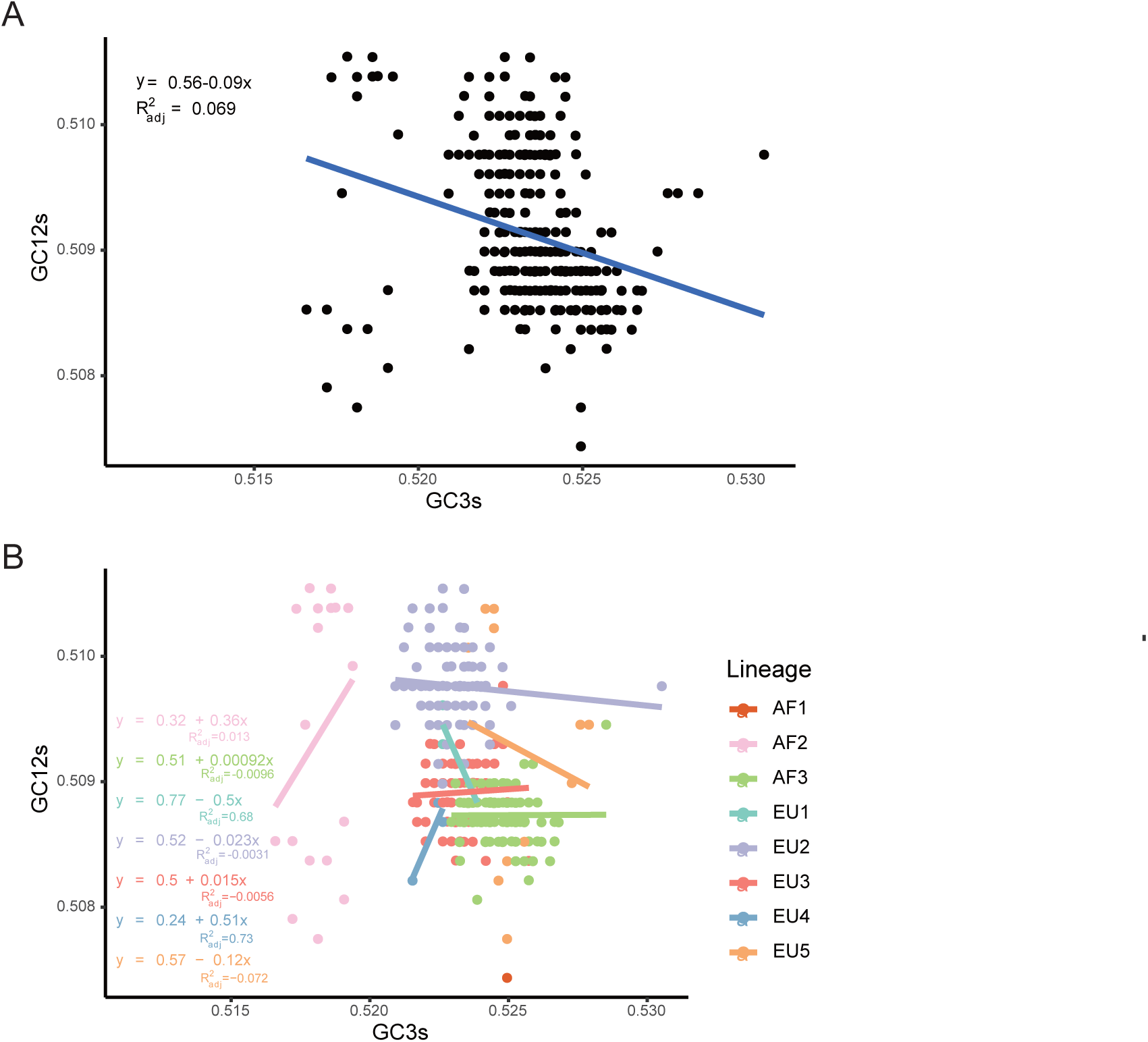
Neutrality analysis of the USUV whole coding sequences for all strains (A) and different lineages (B).

In addition, we performed the neutrality analysis in 10 genes similarly. We found that despite significant correlations between GC12s and GC3s were observed in all genes except the *C* and *NS5*, with relative neutrality ranging from 1.3% (*NS5*) to 23% (*NS1*), mutation pressure was the minor force in all genes, irrespective of lineages (Figure S5A). Taking the lineage information into consideration, all absolute values of regression slopes were less than 0.5 and most of them were close to zero or negative (Figure S5B). The only exception is the *E* genes of the EU4 lineage, but the coefficient between GC12s and GC3s was -0.5 (*P* = 0.55). In a word, although the different degrees of mutation pressure influence in distinct lineages and individual genes, natural selection predominated the evolution of codon usages in USUV.

### Host-specific codon adaptation patterns in USUV

To estimate the relative adaptation of USUV to their hosts and vectors, we performed a CAI analysis. Here we found that the CAI values varied from host to host (Figure 7A). Regarding the whole coding sequence in USUV, the highest CAI values were found in *S. vulgaris* (0.801 ± 0.002), followed by *H. sapiens* (0.796 ± 0.001) and *E. caballuss* (0.773 ± 0.001). The USUV also displayed high CAI values towards the other three NHM hosts. The lowest CAI values were found in *T. merula* (0.508 ±0.002), followed by *P. domesticus* (0.594 ± 0.002) and *Cx. pipiens* pallens (0.618 ±0.001). Except for the pair of two *Culex* species, the CAI values of USUV showed statistical significance in all host pairs (adjusted *P* < 0.05). Taking virus lineages into consideration, we observed that the CAI values of different lineages to the same host varied but a similar pattern was still preserved (Figure 8A). In addition, the CAI values for different hosts varied but relatively conserved patterns were maintained at individual genes across different lineages (Figure 7A).

**Figure 7.**
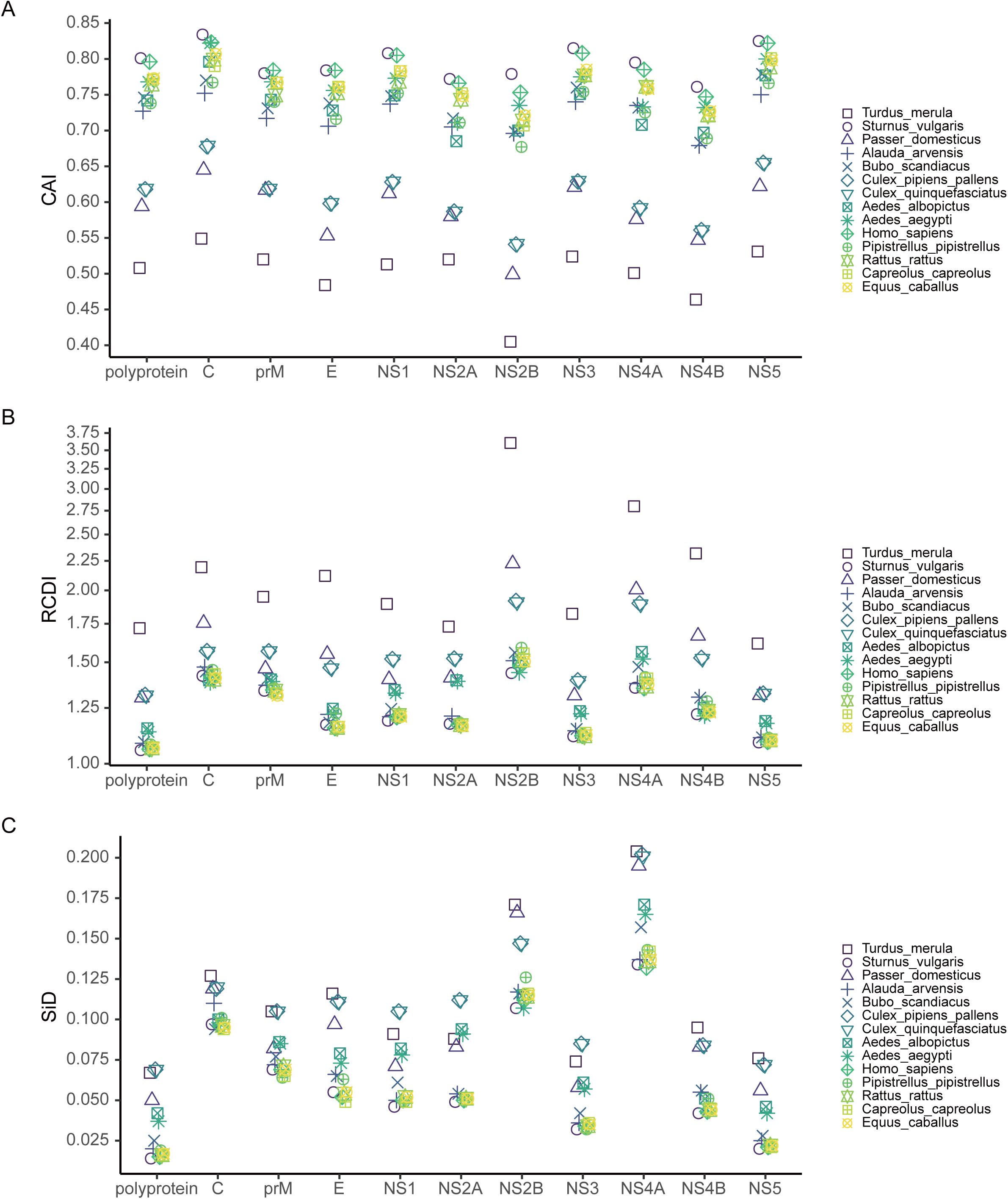
(A) CAI, (B) RCDI, and (C) SiD analysis of the codon usage between USUV coding sequences and their hosts. Different hosts are depicted in distinct shapes and colours.

**Figure 8.**
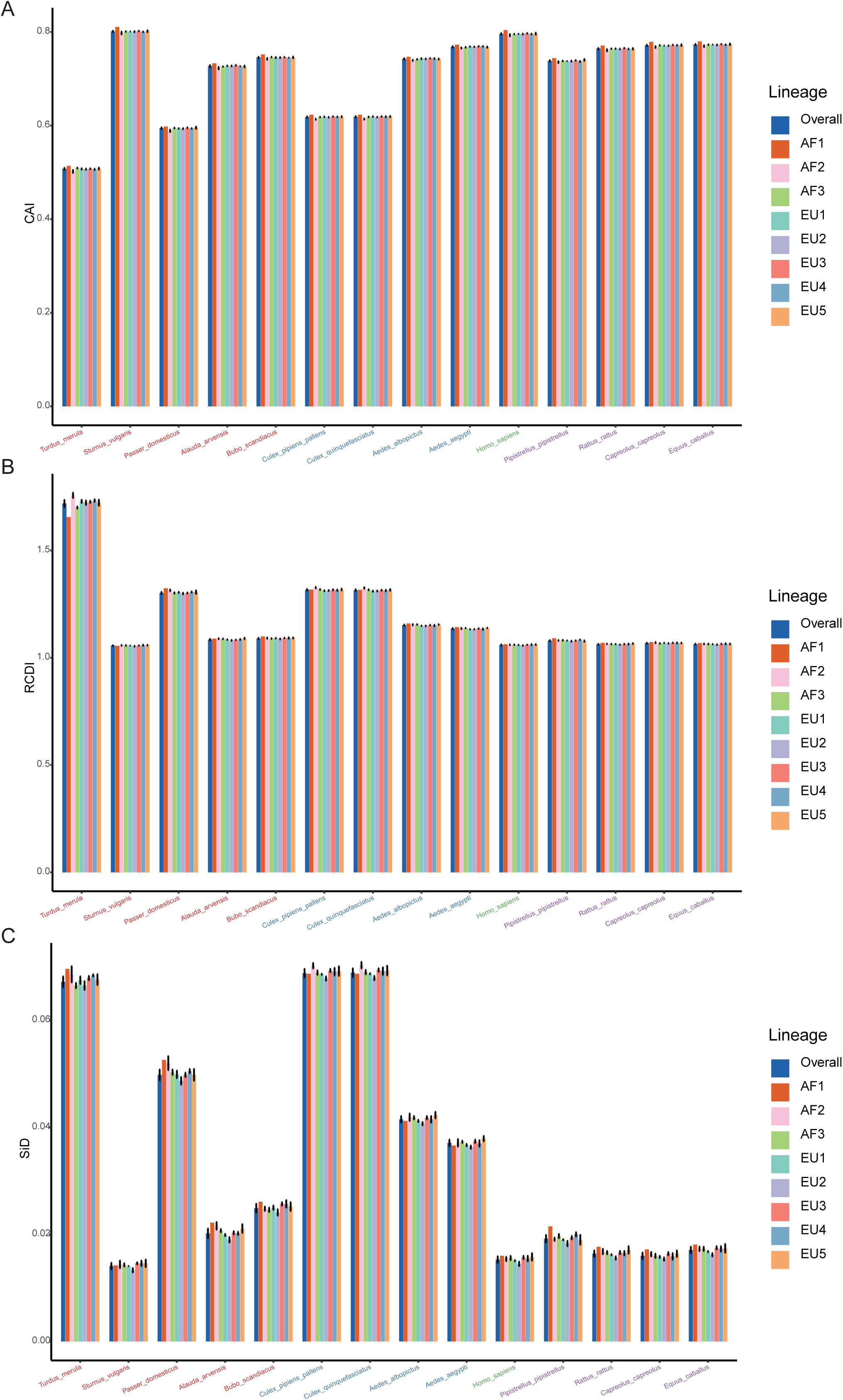
(A) CAI, (B) RCDI, and (C) SiD analysis of the codon usage between the complete coding sequences of the USUV and its hosts. Trends in overall and different lineages are depicted in distinct colours.

### USUV displays the highest codon deoptimization to T. merula

To determine the codon usage deoptimization of the USUV coding sequences with their potential hosts, the RCDI values were inferred. The highest three mean RCDI values were obtained relative to *T. merula* (1.719 ± 0.016), *Cx. pipiens* pallens (1.317 ± 0.004), and *Cx. quinquefasciatus* (1.315 ± 0.004), whereas the lowest three RCDI values were obtained relative to *S. vulgaris* (1.057 ± 0.002), *H. sapiens* (1.060 ±0.002), and *E. caballuss* (1.064 ± 0.002) (Figure 7B). Despite the variation, a similar RCDI values pattern of complete coding sequences in USUV was maintained across all lineages (Figure 8B).

### Cx. quinquefasciatus plays a significantly stronger selection pressure on USUV

To investigate the potential impact of these hosts on the evolution of codon usage patterns of the USUV, the SiD analysis was performed. The results showed that the overall mean SiD value was highest in *Cx. quinquefasciatus* (0.0689 ± 0.0008) versus the complete coding sequences of USUV (Figure 7C). A slightly smaller but comparable (adjusted *P* = 0.77) SiD value was observed in *Cx. pipiens* pallens (0.0688 ± 0.0008). The SiD values in these two hosts were remarkably larger than that in the other hosts simultaneously (adjusted *P* < 0.0001). When considering the lineage classification of the polyprotein sequences, a similar trend remained in all lineages except for the AF1 (Figure 8C), where the highest SiD value was observed in *T*.*merula*, indicating that *Cx. quinquefasciatus* played the strongest influence on the USUV codon usage choices in most of the lineages.

Additionally, the SiD analysis was performed on ten genes of the eight lineages. The mean SiD value for *T. merula* was found to be highest in the *C, E,NS2B, NS4A, NS4B*, and *NS5* genes, while that was found for *Cx. quinquefasciatus* in the *prM, NS1, NS2A*, and *NS3* genes, without consideration for lineages (Figure 7C).There is no significant difference between the SiD values for *Cx. quinquefasciatus* and *Cx. pipiens* pallens in all individual genes. In summary, *Cx. quinquefasciatus* and *T. merula* exerted larger selection pressure on the various genes of different lineages.

## Discussion

In this study, we conducted a comprehensive analysis of the phylogenetic relationships and the codon usage patterns of the USUVs to understand their molecular evolution. Our phylogenetic tree divided the USUV strains into eight lineages. This result is consistent with the previous reports [4,15,34]. The PCA results confirmed the phylogenetic analysis, as the well-defined clusters corresponded to the phylogenetic lineages. This also indicates the USUV has evolved into lineage-specific RSCU patterns, which implies a non-negligible role of evolutionary processes affecting its codon usage.

The genomic composition can greatly affect the CUB [8]. Our data showed that G and A were more abundant in USUV coding sequences. Besides, the RSCU analysis showed that C-end and A-end codons were mostly preferred. These results confirmed a codon bias among the USUV genomes. ENC analysis showed that the overall mean ENC value of all USUV isolates was 55.30, indicating a slightly biased and conserved codon usage. Similar low CUB has also been found in many RNA 346 viruses, such as ZIKV (53.93) [13], JEV (55.30) [35], WNV (53.81), EBOV (55.57) [14], and MARV (ENC, 54.2) [8]. Previous studies suggested lower CUB could reduce the translation resources competition between viruses and their host, which improves viral replication efficiency [8,13]. Therefore, it seems that the low CUB of USUV may have prompted maintaining its circulation in various hosts with different codon usage preferences.

To clarify the factors that influenced the codon usage patterns of USUV, we performed a detailed ENC-GC3s plot, correlation analysis, and neutrality analysis. When the ENC and GC3s values of complete coding sequences of USUV were depicted, we found that all strains were lying below the expected ENC curve, demonstrating that natural selection overall predominated the codon usage of USUV over mutation selection. However, a few contrary phenomena were observed when this analysis was conducted at the level of the individual gene, showing that the effect of mutation pressure was not entirely lacking, especially in some genes such as the *NS2B* and *NS4B*. Correlation analysis reaffirmed the role of natural selection and mutation pressure. Moreover, detailed neutrality analyses demonstrated the dominant role of natural selection in shaping the CUB of USUV, regardless of the lineages and genes. Our results are consistent with the other viruses in the genus *Flavivirus*, such as ZIKV [13] and JEV [35].

The codon usage pattern of the virus is likely affected by its host. Here, we found a mixture of coincidence and antagonism in the codon usage between the USUV and its hosts. This pattern indicated that multiple hosts may have applied selection pressure on the codon usage of the USUV, like ZIKV[13] but different from MARV[8]. Moreover, the CAI and RCDI analysis revealed a disproportionate level of adaptation to its different hosts and vectors, indicating that natural selection exerted pressure on the codon usage of USUVs, although at the variable level from varied hosts. The high adaptation to *H. sapiens* and other mammals suggested the USUV has adjusted its codon usage choice to employ the translation resources more efficiently in mammals, warning of the potential role of these animals in USUV amplification and epidemic. Low adaptation to *T. merula* and *Cx. pipiens* indicated that USUVs have maintained a relatively low translation rate of viral proteins in these hosts, which may be milder harm for these hosts but supports stable survival and spread of progeny viruses. The lowest adaptation to *T. merula* also suggested that it is the most probable primary natural reservoir of USUVs, which is in line with previous reports [1,36].However, our findings are partly inconsistent with Zecchin B *et al* [15], who observed lower CAI values for *S. vulgaris* than *H. sapiens*. This discrepancy might be owing to the different codon usage of *S. vulgaris* used. Yet further investigation is necessary. In addition, as revealed by the SiD analysis, *Cx. quinquefasciatus* have exerted larger selection pressure on the codon usage of 7 of 8 USUV lineages, implying that *Cx. quinquefasciatus* is a potential new favoured vector of USUV. When evaluated in individual genes, the most selection pressure of codon usage of USUV came from *T. merula* and *Cx. quinquefasciatus*, depending on the genes. Accordingly, it makes sense that USUV evolved a lower level of adaptation with its natural reservoir and primary vector than the terminal hosts to facilitate their long-term survival and circulation, as observed in MARV [8] and EBOV [14].

In conclusion, this study reveals a slightly biased and lineage-specific codon usage pattern within USUVs. Mutational pressure, natural selection, and evolutionary processes collectively shaped the codon usage of USUVs. Specifically, natural selection predominated over the other factors. In addition, we found that USUVs have evolved a host-specific adaptation to various hosts and vectors, especially a high fitness to mammals, including humans. The findings of this study improve our insights into the evolution of USUVs that will consolidate future USUV research. Moreover, our results suggest that further epidemiologic monitoring and pathogenicity studies in these high-fitness hosts are particularly required to confront the potential risk of cross-species transmission and outbreak.

## Acknowledgements

This work was supported by the National Natural Science Foundation of China under Grant [number 81830101 and 82072285]; Ministry of Science and Technology of China under Grant [number 2018YFA0902300];

## Disclosure statement

No potential conflict of interest was reported by the author(s).

## Figure Legends

Figure S1. Boxplots of the GC, GC1s, GC2s, and GC3s values of USUV strain in various lineages (A) and isolation hosts (B).

Figure S2. (A) The overall ENC values comparison among the different genes. (B)-(K) show the ENC values of various lineages of the ten genes, respectively.

Figure S3. ENC plots of different genes of the 368 USUV strains. The solid curve represents the expected ENC values when the codon usage was only influenced by the GC3s composition.

Figure S4. Spearman’s correlation analysis among the nucleotide composition, ENC, Aromo, Gravy, and the first two axes of PCA in USUV complete coding sequences. Dark red and blue means positive and negative correlation, respectively. Deeper color darkness means higher correlation. Non-significant (P > 0.05) correlations are not shown.

Figure S5. Neutrality analysis of the USUV genes for all strains (A) and different lineages (B).

## Notes

### Competing Interest Statement

The authors have declared no competing interest.

