## Supplementary figures and images for "A comprehensive analysis of Usutu virus (USUV) genomes revealed lineage-specific codon usage patterns and host adaptation"

### Figure S1

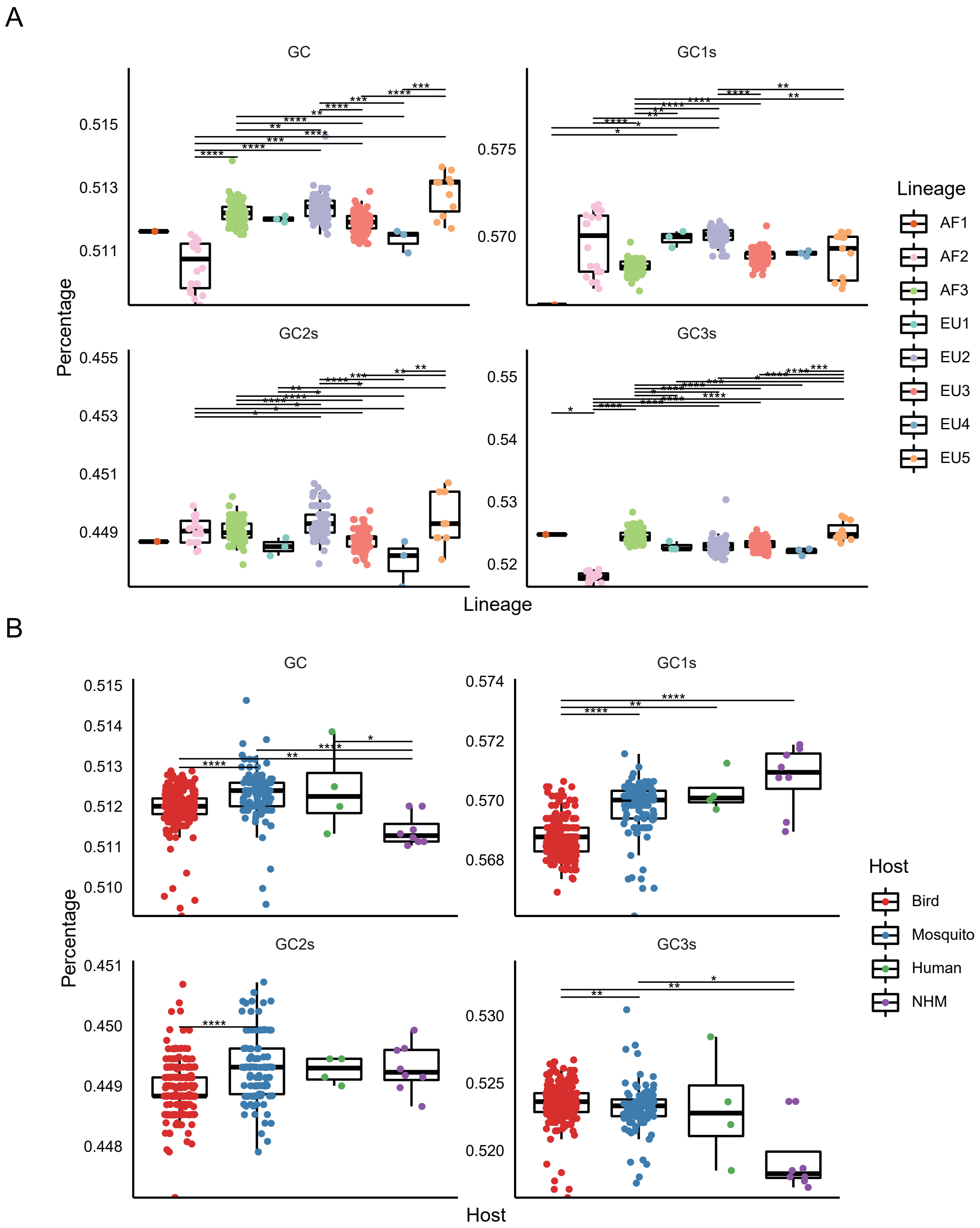

### Figure S2

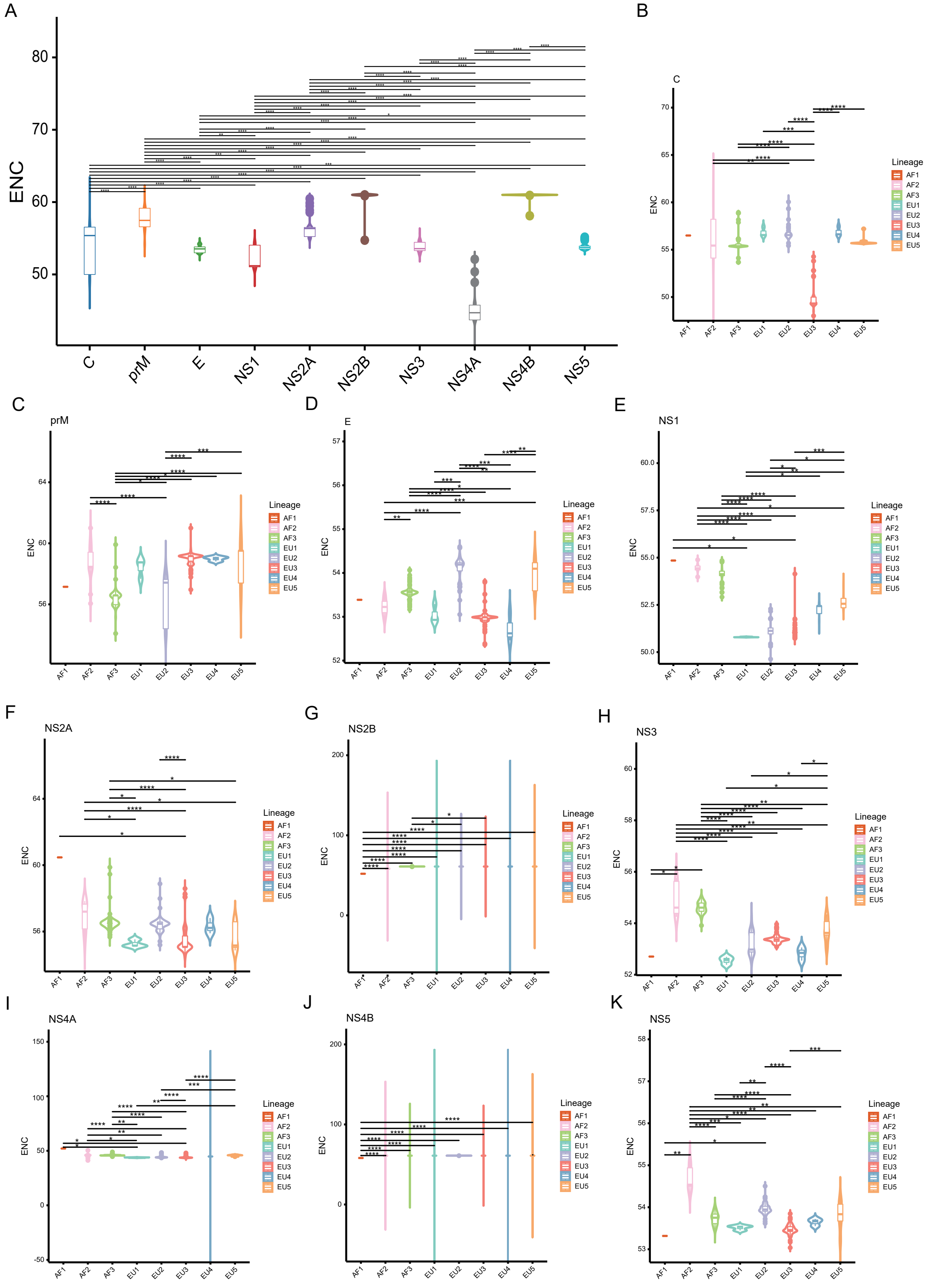

### Figure S3

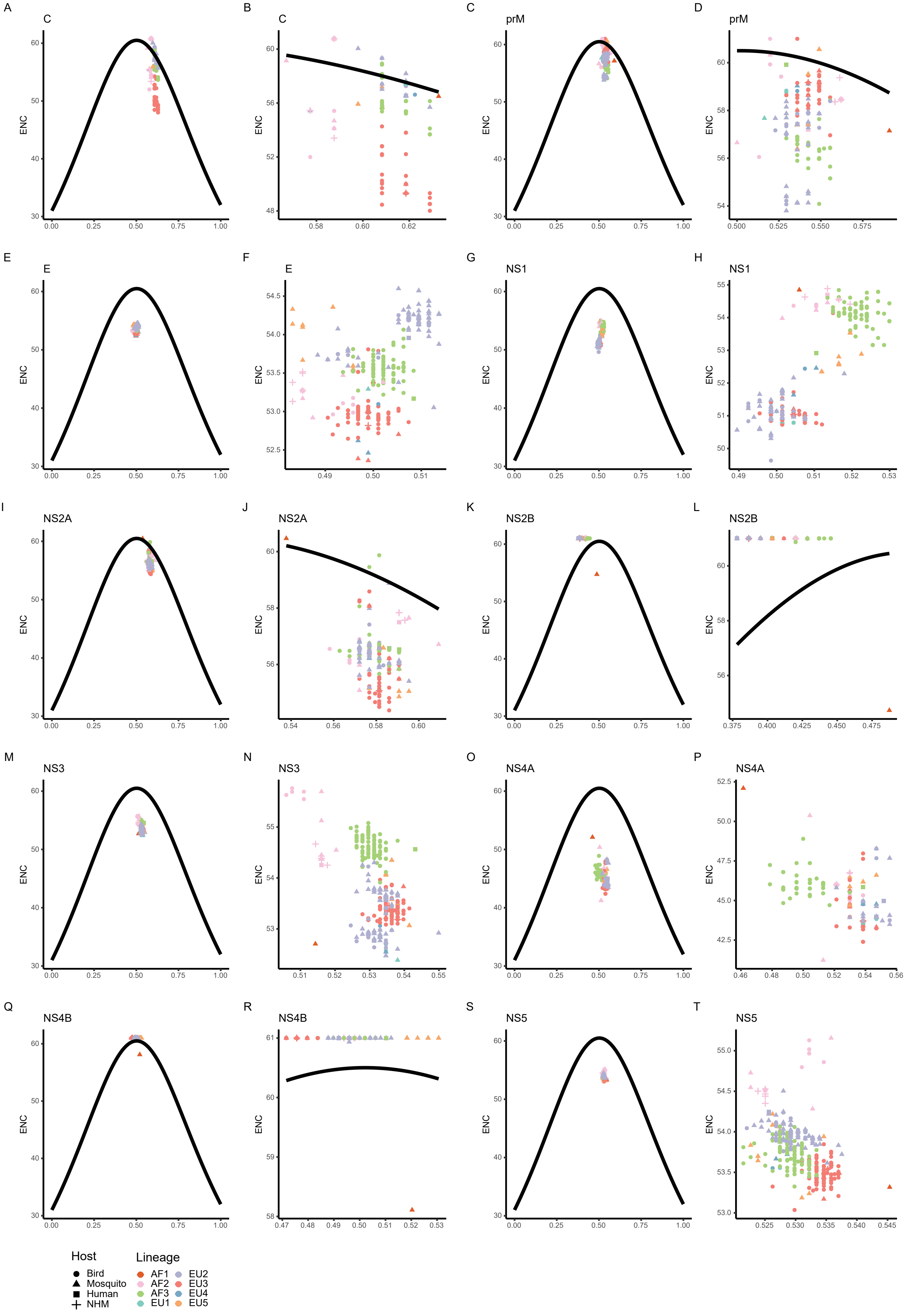

### Figure S4

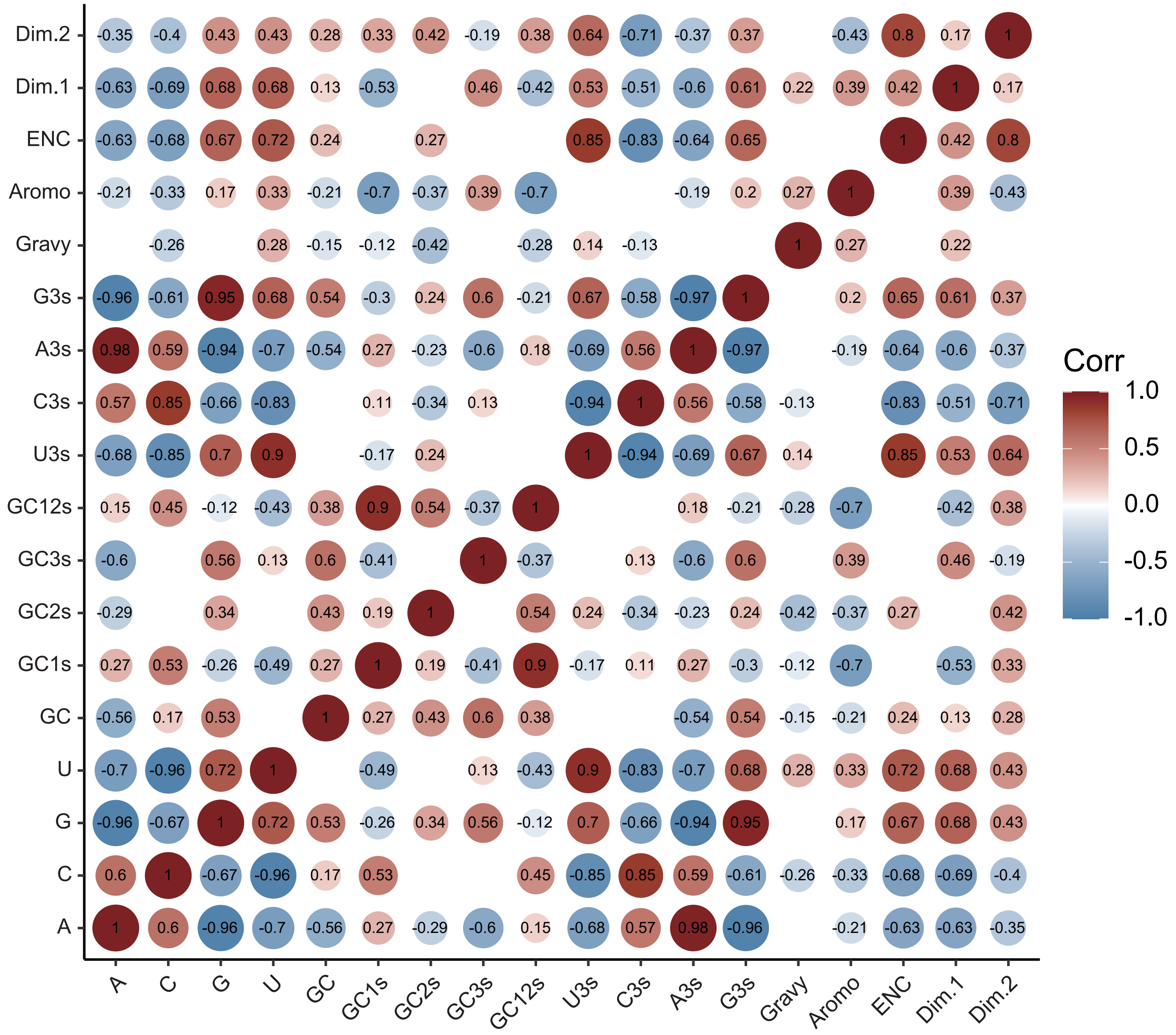

### Figure S5

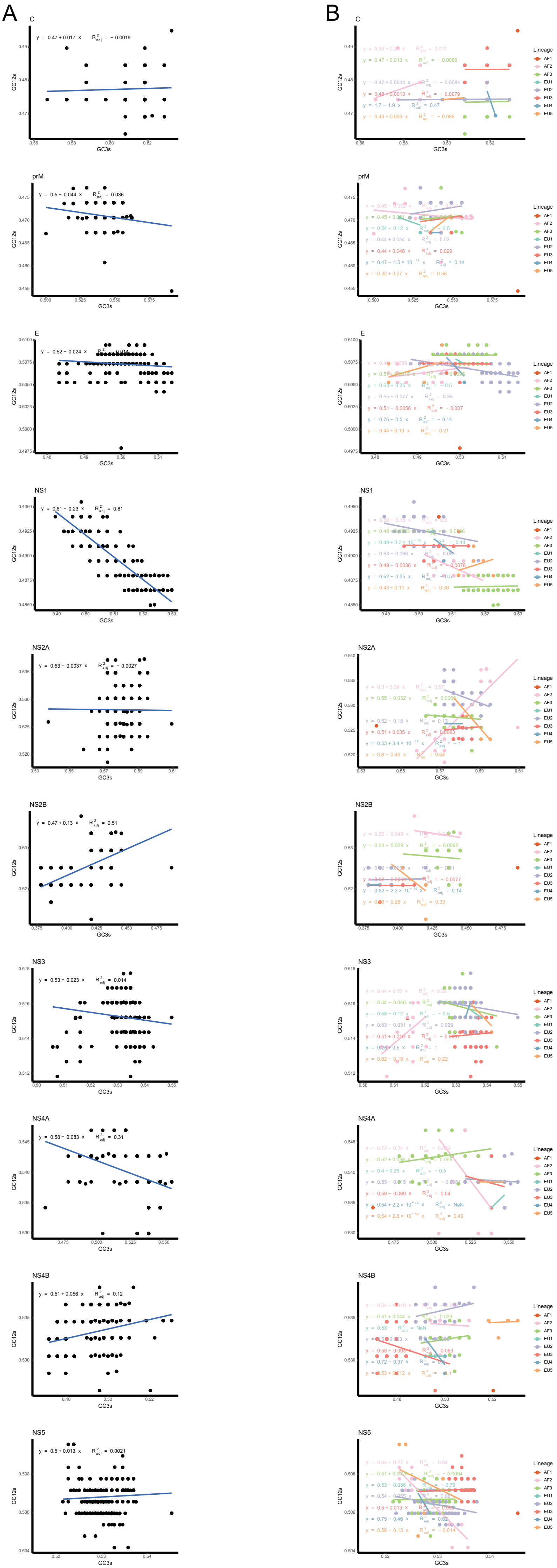
